# Opioid receptor expressing neurons of the central amygdala gate behavioral effects of ketamine in mice

**DOI:** 10.1101/2024.03.03.583196

**Authors:** Matthew B. Pomrenze, Sam Vaillancourt, Pierre Llorach, Daniel Ryskamp Rijsketic, Austen B. Casey, Nicholas Gregory, Juliana S. Salgado, Robert C. Malenka, Boris D. Heifets

## Abstract

Ketamine has anesthetic, analgesic, and antidepressant properties which may involve multiple neuromodulatory systems. In humans, the opioid receptor (OR) antagonist naltrexone blocks the antidepressant effect of ketamine. It is unclear whether naltrexone blocks a direct effect of ketamine at ORs, or whether normal functioning of the OR system is required to realize the full antidepressant effects of treatment. In mice, the effect of ketamine on locomotion, but not analgesia or the forced swim test, was sensitive to naltrexone and was therefore used as a behavioral readout to localize the effect of naltrexone in the brain. We performed whole-brain imaging of cFos expression in ketamine-treated mice, pretreated with naltrexone or vehicle, and identified the central amygdala (CeA) as the area with greatest difference in cFos intensity. CeA neurons expressing both µOR (MOR) and PKCδ were strongly activated by naltrexone but not ketamine, and selectively interrupting MOR function in the CeA either pharmacologically or genetically blocked the locomotor effects of ketamine. These data suggest that MORs expressed in CeA neurons gate behavioral effects of ketamine but are not direct targets of ketamine.

## INTRODUCTION

Ketamine, first approved for human use as a novel anesthetic agent in 1970, has since been recognized as an opioid-sparing analgesic ^1^, a recreational drug with abuse potential ^2^, and most recently, as a rapid-acting antidepressant ^3,4^. The antidepressant effects of ketamine are remarkably rapid, evident within hours of a single infusion, and distinguish ketamine from therapeutics like serotonin-selective reuptake inhibitors which require weeks of daily dosing for efficacy. Ketamine therapy also has some limitations: its profound psychoactive effects require monitoring, single infusions of ketamine improve symptoms of depression for only 1-2 weeks, and long-term ketamine use may be associated with neurological and urological toxicity ^5^. Efforts to improve the safety and durability of ketamine have primarily focused on its antagonism at N-methyl-D-aspartate receptors (NMDARs) in the prefrontal cortex and hippocampus. However, numerous therapeutics based on this mechanism have failed in clinical trials ^6^, strongly suggesting the need for preclinical models that more accurately reflect the antidepressant mechanism of ketamine.

Recent developments have refocused attention on the long-noted interaction between ketamine and opioid receptors (ORs) ^7^, among other receptor targets ^8^. A recent clinical trial found that pre-treating patients with a non-selective OR antagonist, naltrexone, largely blocked the antidepressant effect of ketamine ^9^. Clinical data did not distinguish between a direct effect of ketamine at µ (MOR), δ (DOR) or κ (KOR), versus an indirect effect of naltrexone, for example by disabling endogenous opioid signaling which is known to mediate rapid antidepressant placebo responses ^10^. Subsequent studies in mouse and rat testing the ketamine enantiomers have found *in vitro* and radiographic evidence that both (S)- and (R)-ketamine bind to MORs and KORs with an affinity marginally lower than at NMDARs, and both racemic ketamine and (S)-ketamine produced locomotor activation, locomotor sensitization, and could be self-administered in an OR-dependent fashion ^7,11,12^. Preclinical studies using racemic ketamine have shown that, indeed, this interaction between ketamine and the opioid system is not consistently reflected in conventional assays used to predict antidepressant efficacy, such as the forced swim test (FST), the tail suspension test (TST) and sucrose preference test (SPT) in two stressor models of depression ^13^. Immobility in the FST and TST measured one day post ketamine is unaffected by opioid antagonists ^13^, except potentially at high doses^14^. At shorter timescales, before metabolic clearance of ketamine, studies in unstressed mice show that ketamine effects in the TST and FST can be blocked by naltrexone and a KOR-selective antagonist ^15,16^. Sub-acute effects of ketamine in the FST in congenitally stressed rats, could be blocked by naltrexone, but could not be reproduced by morphine, suggesting a permissive role of the endogenous opioid system ^17^.

To improve our preclinical modeling of the therapeutic effects of racemic ketamine, we systematically screened a range of ketamine-evoked behaviors in mouse for sensitivity to naltrexone. We found that locomotor activation and sensitization, rather than analgesia, the FST, or conditioned place preference, best reflected the ketamine / OR interaction, mediated by MORs. As the bulk of preclinical mechanistic research has focused on brain regions sensitive to NMDAR-like effects of ketamine, we performed an unbiased whole-brain screen for regions reflecting the interaction of ketamine and ORs, by imaging drug-induced expression of the immediate early gene, cFos. This scan, along with fluorescence in situ hybridization, identified neurons of the central amygdala (CeA) which co-express Protein Kinase C δ (PKCδ) and MOR, and were strongly activated by naltrexone rather than ketamine. Intra-CeA microinjection of an MOR antagonist and conditional deletion of MORs from CeA neurons fully blocked ketamine-induced locomotor activation and sensitization.

## RESULTS

### Ketamine-induced hyperlocomotion is opioid receptor-dependent

Ketamine has well described acute and sub-acute behavioral effects in laboratory animals; however, it is unclear which of these behavioral effects best reflect the interaction between ketamine and ORs. We tested for ketamine effects that could be reversed by the broad-spectrum opioid antagonist, naltrexone, in assays of locomotion, analgesia and chronic stress models of depression-like behavior. To assay locomotion, we first pretreated wild-type male mice with naltrexone (NAL; 5 mg/kg, i.p.) or saline (SAL) 30 min before administration of ketamine (KET; 10 mg/kg, i.p.) in an open field test (Fig. 1a). Male mice pretreated with SAL and given KET exhibited the expected increase in locomotor activity. Pretreatment with NAL blocked KET-induced hyperlocomotion (Fig. 1b,c). NAL on its own had no significant effect on locomotion (Fig. 1b,c). We also observed KET-induced hyperlocomotion in female mice, however this effect was not blocked by NAL (Fig. S1a-c), an important sex difference consistent with other reports ^7,12^.

**Figure 1.**
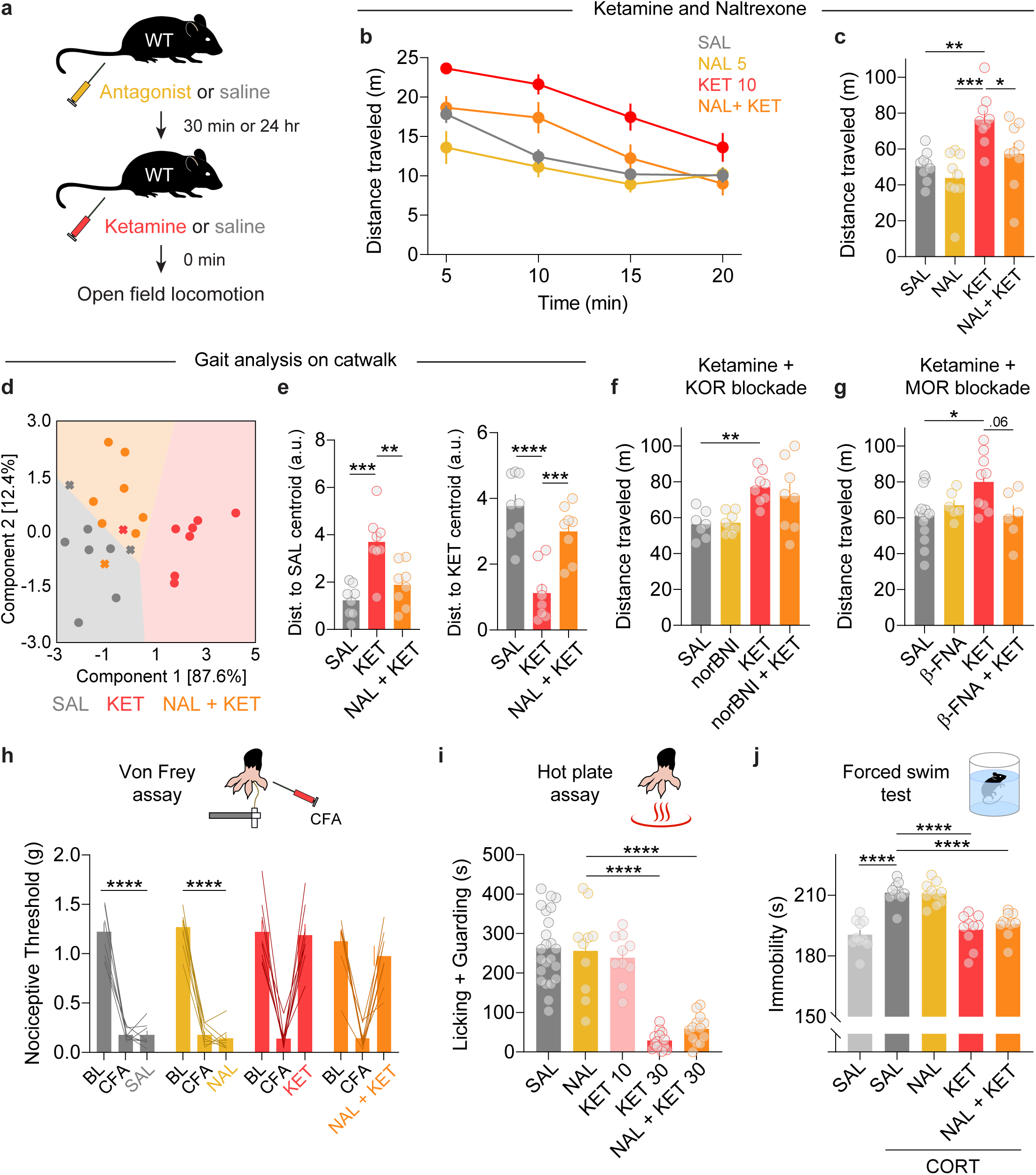
Ketamine locomotor effects require mu opioid receptor activity. a) Schematic of experimental procedure and time points in male mice for panels b-g. b) Time course of locomotor activity after acute SAL (n=9), KET (10 mg/kg; n=9), NAL (5 mg/kg; n=9), or NAL+KET (n=9). c) Summary of total distance traveled in the open field test. KET increased total distance traveled, an effect blocked by NAL. Ordinary one-way ANOVA, F_3,32_ = 8.284, ***p = 0.0003. Tukey’s multiple comparisons test, KET vs SAL **p = 0.002, KET vs NAL ***p = 0.0001, KET vs NAL + KET *p = 0.025. d) Two-component linear discriminant analysis (LDA) of mouse gait after SAL (n=8), KET (10 mg/kg; n=8), or NAL+KET (n=8). See Table S1 for component 1 and 2 feature weights. Colored fields represent LDA decision boundaries. Circles represent accurate classification, X indicates mismatch. A significant drug effect was observed. MANOVA, Pillai’s trace = 0.30, F_2,21_ = 4.57, p = 0.022. e) Distance (arbitrary units [a.u.]) of all groups to either the SAL (*left*) or KET (*right)* centroid. KET subjects are a significant distance from both SAL and NAL+KET centroids. ANOVA, SAL centroid F_2,21_ = 13.74, p = 0.0002; Tukey’s multiple comparisons test, NAL+KET vs SAL p = 0.39, KET vs SAL *** p = 0.0001, KET vs. NAL+KET ** p = 0.0035. ANOVA, KET centroid F_2,21_ = 19.31, p < 0.0001; Tukey’s multiple comparisons test, SAL vs KET **** p < 0.0001, NAL+KET vs KET *** p = 0.0009, SAL vs NAL+KET p = 0.2. f) KET-induced locomotion was not blocked by KOR antagonist norBNI (given 24 hours prior). SAL, n=7; norBNI, n=8; norBNI+KET, n=8. Ordinary one-way ANOVA, F_3,27_ = 5.866, **p = 0.0032. Tukey’s multiple comparisons test, KET vs SAL **p = 0.0073, KET vs norBNI **p = 0.0076, KET vs norBNI + KET p = 0.77. g) KET-induced locomotion was attenuated by MORs antagonist β-FNA (given 24 hours prior). SAL, n=15; β-FNA, n=6; β-FNA+KET, n=6. Ordinary one-way ANOVA, F_3,32_ = 3.971, *p = 0.016. Tukey’s multiple comparisons test, KET vs SAL **p = 0.014, KET vs β-FNA p = 0.29, KET vs β-FNA + Ket p = 0.061. h) Enhancement of nociceptive thresholds from acute KET after inflammatory pain induction with CFA (24 hrs prior) was not NAL-sensitive (von Frey assay). Thresholds were tested at pre-CFA baseline (BL). SAL, n=8; NAL, n=8; KET, n=8; NAL+KET, n=8. 2-way RM ANOVA, interaction of pain state and drug condition, F_6,56_ = 17.67, ****p < 0.0001. Tukey’s multiple comparisons test, SAL – BL vs CFA ****p < 0.0001, BL vs SAL ****p < 0.0001, CFA vs SAL p > 0.99. Nal – BL vs CFA ****p < 0.0001, BL vs NAL ****p < 0.0001, CFA vs NAL p = 0.75. KET – BL vs CFA ***p = 0.0002, BL vs KET p = 0.9824, CFA vs KET ***p = 0.0002. NAL + KET – BL vs CFA ****p < 0.0001, BL vs KET p = 0.4812, CFA vs KET ***p = 0.0006. i) KET-diminished non-reflexive (affective) pain responses were not NAL-sensitive (hot plate assay). SAL, n=23; NAL, n=10; KET 10, n=10; KET 30, n=14; NAL+KET30, n=13. Ordinary one-way ANOVA, F_4,65_ = 36.61, ****p < 0.0001. Tukey’s multiple comparisons test, SAL vs KET 30 ****p < 0.0001, SAL vs NAL + KET 30 ****p < 0.0001, KET 10 vs KET 30 ****p < 0.0001, KET 10 vs NAL + KET 30 ****p < 0.0001. j) KET (30 mg/kg 24 hours before testing) reduced immobility in the FST in mice after chronic CORT treatment. NAL did not reverse this effect. SAL, n=10; CORT: SAL, n=10; NAL, n=10; KET, n=10; NAL+KET, n=10. One way ANOVA, F_4,45_ = 22.64, ****p < 0.0001. Tukey’s multiple comparisons test, SAL vs SAL CORT ****p < 0.0001, SAL CORT vs KET ****p < 0.0001, SAL CORT vs NAL + KET ****p < 0.0001.

To determine whether OR involvement is unique to hyperlocomotion induced by KET versus other stimulants, we tested whether NAL could block cocaine (COC)-induced hyperlocomotion. We found no difference in distance traveled between COC versus NAL+COC treated groups (Fig. S1d). Furthermore, we found distinct effects of COC and KET on gait, measured while mice traversed a ‘catwalk’ apparatus (Fig. S1e,f; Table S1). Linear discriminant analysis of dimensionality-reduced gait parameters effectively separated COC, KET, and SAL-treated mice, suggesting different mechanisms underlying the locomotor effects of these drugs. A similar gait analysis showed that NAL prevented KET’s effects on gait (Fig. 1d,e; Table S2), consistent with blockade of hyperlocomotion. NAL is not selective for OR subtypes. We therefore tested KET-induced hyperlocomotion for sensitivity to a selective KOR antagonist (norbinaltorphimine [norBNI]), and a MOR antagonist (beta-funaltrexamine [β-FNA]). To accommodate the slow pharmacokinetics of these antagonists ^18,19^, we tested KET in the open field 24 hr after pretreatment with either antagonist. While norBNI failed to significantly block KET’s locomotor effect (Fig. 1f), β-FNA effectively reduced KET-induced hyperlocomotion to the control level (Fig. 1g).

KET is a well-known analgesic. We therefore tested whether NAL could block KET analgesia in an inflammatory pain model, using an injection of Complete Freund’s Adjuvant (CFA) in the right hindpaw. As expected, KET had an analgesic effect on measures of both reflexive analgesia (paw withdrawal threshold in the Von Frey assay, Fig. 1h) and non-reflexive (‘affective’) analgesia (licking and guarding behavior in the hot plate assay, Fig. 1i). Pretreatment with NAL had no effect on either form of KET-induced analgesia.

KET has widely reported effects on the FST, a measure of depression-like behavior in which reduced immobility is interpreted as an active coping response ^4^. We tested both unstressed mice and mice treated with 10 days of 1% corticosterone (CORT) in their drinking water, the latter which increased immobility in the FST (Fig. 1j). We were able to elicit a reduction in immobility in both stressed and unstressed mice, 24 hr after KET treatment, using a higher dose of KET (30 mg/kg, i.p.) than required for locomotor and analgesic effects (Fig. 1j; Fig. S1g). Mice pretreated with NAL 30 min prior to KET treatment were indistinguishable from mice treated with KET alone regardless of stress exposure. We were unable to elicit KET effects in the TST 24 hrs following treatment, or in a KET conditioned place preference assay (20 mg/kg, i.p.) (Fig. S1h,i).

Altogether, these data suggest that the MOR-dependent locomotor stimulant effect of KET best reflects the finding in humans that KET’s antidepressant effect is OR-dependent. Engagement of ORs is not required for KET’s effects on pain or in assays widely used to study antidepressant-like effects of KET in mice.

### Brain-wide activity mapping localizes a KET / OR interaction to the central amygdala

Most investigations of KET’s antidepressant mechanisms in mice have focused on structural and functional changes in prefrontal cortex and hippocampus ^20,21^, interpreted with behavioral assays like the FST. To identify the neural substrates mediating the interaction between KET and MORs, read out by KET’s locomotor effect, we quantified neural activity, indexed by cFos expression, across iDISCO+ cleared mouse brains using UN-biased high-Resolution Analysis and Validation of Ensembles using Light sheet images (UNRAVEL) ^22^. We performed activity mapping in Fos^2A-iCreER^(TRAP2);Ai14 mice to obtain two independently quantified read-outs of whole brain activity, cFos and tdTomato (Fig. S2a). To obtain both measures in each mouse, we adopted a crossover treatment strategy (Fig. S2b). After an injection of 4-hydroxytamoxifen (4-OHT, 50 mg/kg) to activate Cre^ER^ and tdTomato expression, mice were pretreated with SAL or NAL (5 mg/kg i.p.), followed 30 min later by KET (10 mg/kg, i.p.). One week later, to ensure adequate washout of any residual behavioral effects ^21,23^, mice were crossed over to the opposite treatment and perfused 90 min later to capture peak cFos expression. After fixation, brain hemispheres were stained for cFos or tdTomato, cleared using iDISCO+, and image volumes were obtained via light sheet fluorescence microscopy (LSFM) (Fig. 2a,b). Our primary outcome was a whole-brain voxel-based comparison of cFos expression in SAL+KET versus NAL+KET image volumes. Clustered hotspots were validated with density measurements of segmented cFos+ cells in the same areas (Fig. 2c-f). Validated hotspots obtained from cFos mapping were verified by independent quantification of tdTomato cell densities within cFos-defined anatomical clusters (Fig. S2).

**Figure 2.**
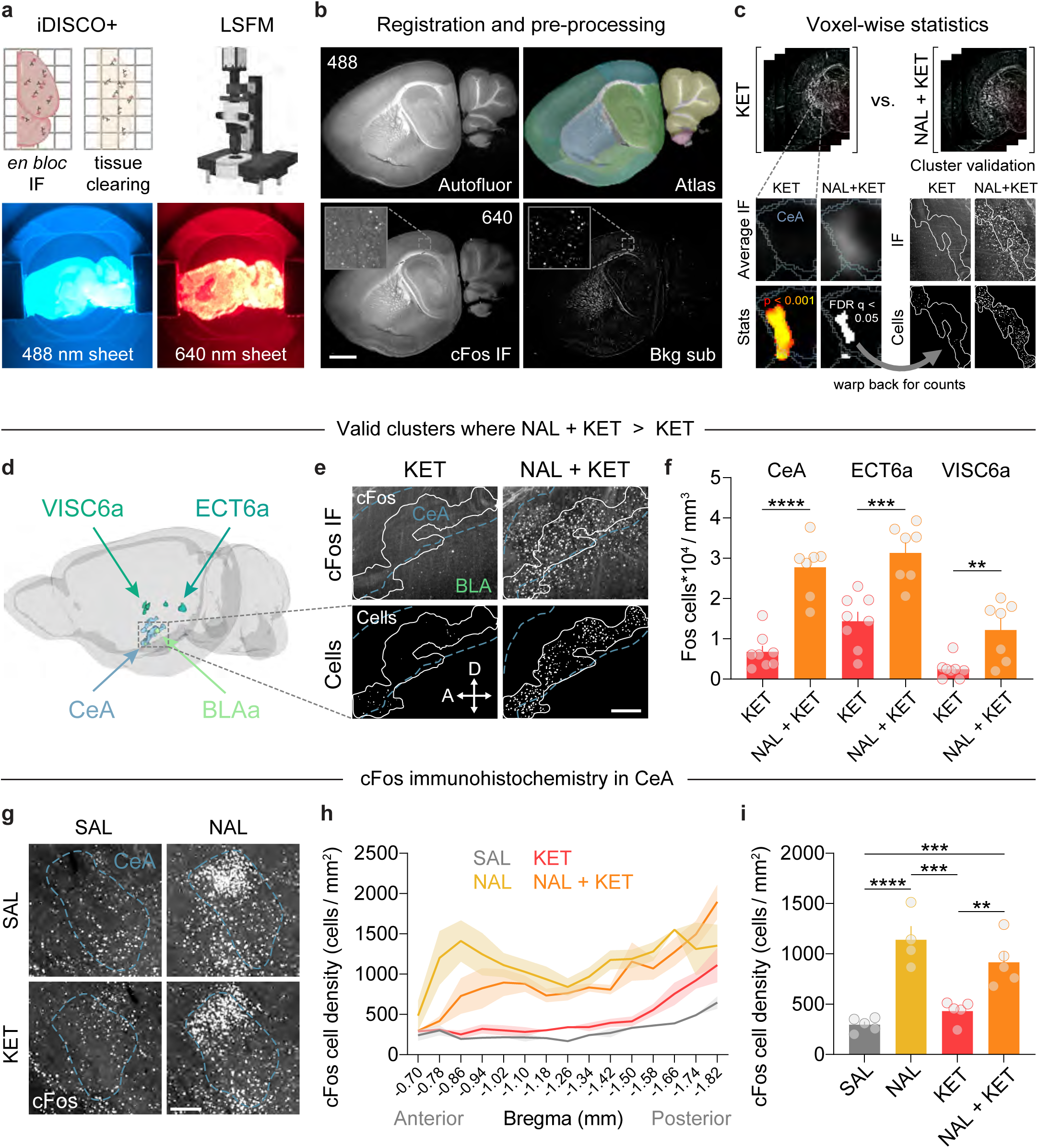
Whole brain mapping localizes a KET / OR interaction to the central amygdala. a) Mice were injected with NAL (5 mg/kg) or SAL 30 min prior to injecting KET (10 mg/kg). Brains were then immunolabeled for cFos, cleared with iDISCO+, and imaged in 3D with a light sheet microscope. 488 nm light excited autofluorescence, whereas 640 nm light evoked fluorescence from cFos immunoreactivity. b) Autofluorescence was registered with an iDISCO-specific averaged template brain in alignment with the Allen brain atlas (CCFv3). cFos immunofluorescence (IF) was rolling ball background subtracted and warped to atlas space using transforms from registration. Scale = 1 mm. c) cFos-IF images in atlas space were z-scored and smoothed (50 µm kernel). Increased cFos-IF in the central amygdala (CeA) from NAL was notable when comparing averaged images. This effect was confirmed by voxel-wise nonparametric permutation testing and cluster correction (FDR q < 0.05). To validate differences between groups, clusters of significant voxels were warped to full resolution tissue space for cell density measurements using a segmentation of cFos^+^ cells from Ilastik. d) Sagittal view of valid clusters in a 3D brain where cFos-IF was greater with NAL+KET (n=7) than KET alone (n=8). Abbreviations: CeA = central amygdala; ECT6a = Ectorhinal area, layer 6a; VISC6a = visceral area, layer 6a; BLAa = anterior basolateral amygdala. See Table S3 for cluster volumes, locations, and regional compositions. e) Example of cFos-IF and segmentation of cFos^+^ cells in the CeA cluster (white outline) for SAL+KET and NAL+KET groups shown as maximum intensity projections. CeA boundary lines were guided by a median intensity projection of atlas labels. Abbreviations: A = anterior; D = dorsal. Scale = 200 µm. f) NAL increased segmented cFos^+^ cell densities in 3 of 5 clusters. The predominant region of each valid cluster is indicated. *Post hoc* unpaired, two-tailed t-tests by region: CeA, t_13_ = 7.10, ****p < 0.0001; ECT6a, t_13_ = 4.80, ***p = 0.0003; VISC6a, t_13_ = 3.54, **p = 0.0036. g) Mice in a separate cohort were injected as before to quantify the density of cFos^+^ cells in the CeA in coronal brain slices. Examples of cFos-IF are shown for each group. Scale = 200 µm. h) Plot of cFos^+^ cell densities in the CeA across its anterior to posterior axis. i) Quantification of cFos^+^ cell densities in the CeA. Ordinary one-way ANOVA, F_3,15_ = 20.94, ****p < 0.0001. Tukey’s multiple comparisons test, Sal vs Nal ****p < 0.0001, SAL vs KET p = 0.67, SAL vs NAL + KET ***p = 0.005, NAL vs KET ***p = 0.0002, KET vs NAL + KET **p = 0.0045, NAL vs NAL + KET p = 0.31.

Autofluorescence (488 nm excitation) of each brain was registered to an LSFM-specific Allen brain atlas template ^24^. Immunofluorescent staining (640 nm excitation) was rolling ball background subtracted and z-scored across hemispheres. We performed a voxel-wise unpaired t-test between cFos-labelled hemispheres from mice treated with SAL+KET versus NAL+KET; statistical contrasts underwent False Discovery Rate (FDR) correction (q < 0.05) to identify clusters of significant voxels. To ensure that detected clusters represented valid differences in the density of recently activated brain cells, we warped each cluster to full resolution tissue space, quantified the density of segmented cFos^+^ cells, and performed *post hoc* t-tests.

Our voxel-wise analysis identified five clusters surviving FDR correction, with greater cFos immunofluorescence in the NAL+KET group compared to the SAL+KET group. Three of these clusters remained significant after comparing segmented cFos^+^ cell densities between groups. The most substantially altered cluster of neural activity was primarily localized to the capsular region of the central amygdala (CeA) (Fig. 2d-f; Table S3). We also detected differences in layer 6a of the ectorhinal and visceral areas (ECT6a and VISC6a, respectively) (Fig. 2d,f). One cluster, located in the tuberal nucleus of the hypothalamus, was associated with decreased activity following NAL+KET compared to SAL+KET, but was not significant in a *post hoc* unpaired t-test (p = 0.63). These results were reproduced through quantification of tdTomato^+^ cells in the same clusters, albeit with weaker effect sizes (Fig. S2d), consistent with the reported under sampling of cFos cells in TRAP2 mice ^22,25,26^.

We were particularly intrigued by the CeA, as it is enriched with MORs and plays a crucial role in both affective behaviors and freezing associated with fear states ^27–31^. To determine if NAL independently increases activity in the CeA, we performed a detailed analysis of cFos cells in brain slices spanning the entire antero-posterior axis of the CeA with confocal microscopy (Fig. 2g-i). We quantified cFos^+^ cell counts in four conditions: SAL, NAL, SAL+KET, NAL+KET. Whereas KET moderately increased the abundance of cFos cells, particularly in the posterior portions of the CeA, the obviously dominant effect was the enhanced cFos count in the NAL group, evident at virtually every AP position. In comparison to SAL conditions, cFos^+^ cell counts in the NAL+KET groups were elevated to a similar extent as NAL alone, and both NAL and NAL+KET groups contained higher cFos^+^ cell counts compared to SAL+KET.

Our behavioral results and imaging data suggest that disinhibition of opioidergic CeA neurons by NAL is sufficient to block KET-induced hyperlocomotion. Sub-anesthetic KET does not appear to have a major direct effect on CeA activity. If MOR-expressing neurons of the CeA, activated by NAL, effectively gate KET-induced hyperlocomotion, we predicted that: 1) the CeA ensemble activated by NAL expresses MORs; and 2) blocking MORs in CeA either pharmacologically or through genetic manipulation prevents KET-induced hyperlocomotion.

### MORs in the CeA are critical to ketamine’s locomotor effects

To determine the molecular phenotype of NAL-activated neurons in the CeA, we assessed the overlap between cFos^+^ and MOR^+^ neurons in major cell groups of the CeA. Two major and distinct groups that play critical roles in fear learning and anxiety states express PKCδ or somatostatin (SST) ^28,32,33^. To visualize whether the ensemble of activated cells in the CeA expresses MORs, PKCδ, or SST, we performed fluorescence *in situ* hybridization (FISH) on CeA sections harvested 30 min after an injection of NAL (Fig. 3a). We observed the expected *Fos* ensemble in the capsular CeA and detected *Oprm1* (gene that encodes MOR) in roughly 80% of *Fos* cells (Fig. 3b-d). When probing for cell markers, we found a striking prevalence of *Fos* positive neurons among the *Pkcd*, but not the *Sst* expressing CeA neurons (Fig. 3b-d). The majority of *Fos* cells expressing *Oprm1* also expressed *Pkcd*. These FISH data suggest that CeA cells co-expressing PKCδ and MOR might gate OR-sensitive locomotor effects of ketamine.

**Figure 3.**
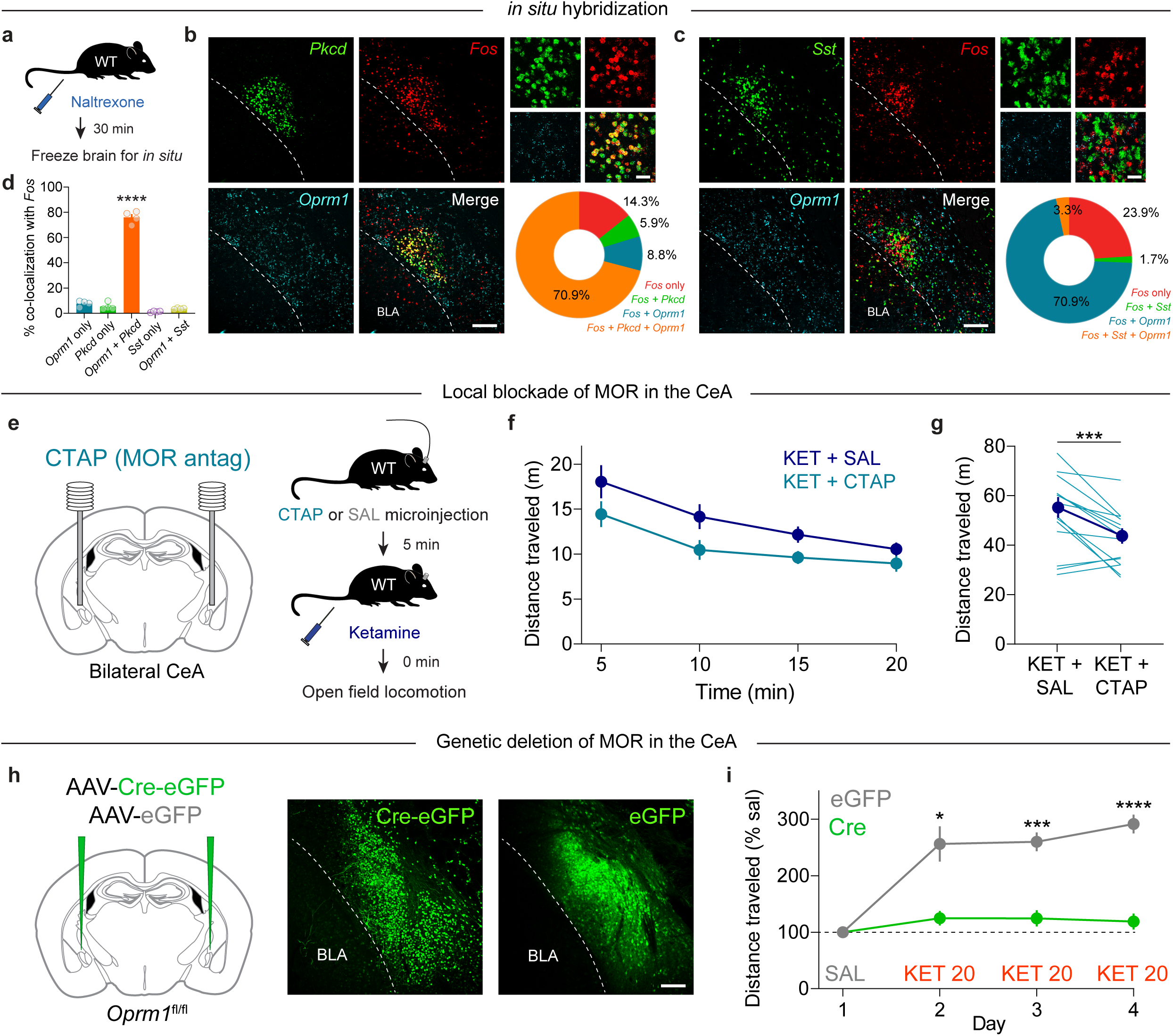
MORs in CeA PKCd neurons are critical to ketamine’s locomotor effects. a) Schematic of experimental design for fluorescence in situ hybridization. b) Representative images and pie chart showing high degree of overlap between *Pkcd*, *Fos*, and *Oprm1* in the CeA after Nal administration. Scale = 100 µm. High magnification image, scale = 10 µm. c) Representative images and pie chart showing low degree of overlap of *Sst* with *Fos* and *Oprm1* in the CeA after NAL administration. Scale = 100 µm. High magnification image, scale = 10 µm. d) Quantification of co-localization of marker genes with *Fos* in the CeA. The majority of *Fos*^+^ cells were labeled with *Oprm1* and *Pkcd*. N=4 wild-type mice. Ordinary one-way ANOVA, F_4,15_ = 526.0, ****p < 0.0001. e) Left, setup of MOR antagonist microinjection into CeA. Right, experimental procedure. f) Time course of locomotor activity after administration of KET (10 mg/kg) and CTAP microinjection (300 ng in 500 nL / hemisphere). KET+SAL, n=15; KET+CTAP, n=15; crossover design. g) CTAP significantly reduced ketamine-induced locomotor behavior in the open field. Paired, two-tailed t-test, t_14_ = 4.19, ***p = 0.0009. h) *Left*, injection of AAV-Cre-eGFP into the CeA of *Oprm1*^fl/fl^ mice for genetic deletion of MOR. *Right*, representative images of Cre-eGFP and eGFP expression in the CeA. Scale = 100 µm. i) Time course of locomotor sensitization to KET (20 mg/kg) in mice with genetic deletion of MOR in the CeA (Cre, n=14) or a control group (eGFP, n=5), normalized to Day 1 (SAL). 2-way RM ANOVA, interaction of *Oprm1* cKO and drug condition, F_3,51_ = 19.62, ****p < 0.0001. Sidak’s multiple comparisons test, eGFP vs Cre: Day 2 *p = 0.040, Day 3 ***p = 0.0003, Day 3 ****p < 0.0001.

To determine the importance of MORs in the CeA for ketamine’s effects on locomotion, we implanted bilateral guide cannulas targeting the CeA into wild-type male mice (Fig. 3e). Mice were locally microinjected with the fast-acting, reversible MOR antagonist CTAP or SAL 5 min before KET (10 mg/kg, i.p.) and tested in the open field (Fig. 3e). These same mice were then microinjected again one week later in a crossover design. KET locomotion was reduced by intra-CeA CTAP infusion (Fig. 3f,g), indicating a role for MOR activation in the CeA for this behavior. A subsequent test of CTAP alone showed no difference compared to SAL controls on locomotion (Fig. S3).

Given the relatively modest effect of intra-CeA administration of an MOR antagonist, we tested a more robust behavioral paradigm, locomotor sensitization, based on recent data that ketamine and (*S*)-ketamine-induced locomotor sensitization is NAL-sensitive ^7,12^. We found that 20 mg/kg KET (i.p.), administered daily for three days, produced locomotor sensitization in mice (Fig. 3h,i). To avoid multiple rounds of intracerebral injection, and to account for the inability of drug infusion to distinguish between MORs expressed in local CeA cells versus axon terminals, we used a genetic approach. We injected floxed MOR (*Oprm1*^fl/fl^) mice with AAV-Cre-eGFP to genetically delete *Oprm1* selectively in the CeA cells (Fig. 3h). In *Oprm1*^fl/fl^ mice injected with a control virus (AAV-eGFP), KET locomotor sensitization was robust. In contrast, deletion of *Oprm1* in the CeA with AAV-Cre-eGFP disrupted KET’s locomotor effects (Fig. 3i,j; Fig. S3) at all time points. Together, these data demonstrate that MORs in CeA neurons are necessary for the locomotor stimulating effects of KET.

## DISCUSSION

### Summary

Ketamine is an antidepressant drug with pharmacology and behavioral effects unlike conventional antidepressants and is a promising lead compound for novel therapeutics. However, improving on the serendipity of ketamine has proceeded at a surprisingly slow pace, compared to other mechanism-informed drug development efforts ^34,35^. Major obstacles include the continued use of preclinical behavioral models that do not predict antidepressant efficacy in humans ^36–38^, and a limited set of in-human mechanistic experiments to validate preclinical model predictions ^9,39,40^.

This preclinical investigation stems from recent in-human findings that NAL, a nonselective opioid antagonist, blocks the antidepressant effect of KET ^9^. We screened several KET-evoked behaviors in mice for opioid receptor sensitivity, finding that locomotor activation and sensitization uniquely and robustly reflect the mechanism observed in humans. Using these simple behavioral assays and selective OR antagonists, we narrowed the effect of NAL to antagonism at MORs, and not KORs. MORs are broadly expressed in the brain, therefore we undertook an unbiased approach to identify regions that may mediate the KET / MOR interaction using drug-evoked cFos expression as a surrogate marker for differential neural activity. This brain-wide search revealed that PKCδ - and MOR-expressing neurons of the CeA capsular region showed the greatest differences between SAL+KET versus NAL+KET treatments. Follow-up experiments where we selectively blocked the locomotor activating effects of KET by inactivating MORs in the CeA, either pharmacologically or through conditional KO, fully validated our localization strategy.

MORs may be a direct target for KET; alternatively, a functioning opioid signaling system could be a prerequisite (i.e. “permissive”) for KET to exert its effect. Our finding strongly suggests the latter: while ketamine only marginally increased CeA cFos expression, NAL dramatically enhanced CeA cFos in the same cells. In effect, locking CeA neurons in the “on” state with NAL appears to prevent KET from producing locomotor effects. In this model, KET-induced locomotion, perhaps via NMDAR antagonism and/or striatal dopamine release ^23,41^, is gated by MOR-dependent activity localized to PKCδ CeA neurons.

### Modeling opioid receptor dependent effects of ketamine in rodents

The simple assays in our behavioral screen represent several commonly described effects of KET on rodent behavior, assembled to establish which ones correspond most closely with findings in humans. Hyperlocomotion and sensitization in adult male C57BL/6 mice showed the strongest opioid-dependence. Hyperlocomotion does not have the same face validity as a model of depression-like behavior compared to assays of passive coping behavior, like FST and TST, or anhedonia, like SPT. Rather, drug-evoked hyperlocomotion and sensitization are strongly associated with a drug’s abuse liability in rodents and humans ^42^, including opioids ^43^. KET-induced hyperlocomotion has been reproduced across labs for more than 40 years ^7,11,16,23,44–46^, however few studies have directly tested the role of ORs in this behavior ^7,12,45^.

The timing of measurement, animal sex, and enantiomeric composition of KET are critical factors for reproducing OR-dependent hyperlocomotion. We found that the hyperlocomotion evident within first 20 minutes after i.p. injection of low dose KET (10 mg/kg) is most sensitive to OR antagonists, consistent with similar findings with (S)-ketamine in mice ^7^ and racemic KET in rat ^12^, and likely accounting for an early report finding no effect of naloxone on KET-induced hyperlocomotion at later timepoints ^45^. Interestingly, while we and others find that male and female rodents show hyperlocomotor responses to KET ^12,23^, only the male response is blocked by NAL, suggesting that the mouse model that best reflects the human physiology is quite specific, leaving open the question of how to investigate human sex-dependent responses to KET in rodent models. Finally, while (S)-ketamine appears to support a range of behaviors related to abuse liability (i.v. CPP, self-administration, locomotor sensitization) ^7,11^ with the latter two blocked by NAL-it is unclear whether racemic KET has the same features. We were unable to demonstrate a racemic KET CPP, consistent with recent findings that racemic KET has relatively low abuse liability in mice ^23^.

In examining other KET-evoked behaviors, our data consistently parallel human findings. The interaction between ketamine and the opioid system has been extensively documented in the context of analgesia. In humans and laboratory animals, KET substantially potentiates the analgesic effect of opioids, and reduces both opioid tolerance and opioid-induced hyperalgesia ^47–49^, all effects that may contribute to a prolonged reduction in postoperative opioid use, even when KET is delivered during surgical anesthesia ^1^. However, the analgesia produced by KET itself at sub-anesthetic doses is *not* blocked by opioid receptor antagonists in humans ^50^, reflected in our findings in mice using a dose of KET (10 mg/kg i.p.) that is sub-dissociative ^51,52^, and consistent with previous work ^45,53,54^. At high doses (>100 mg/kg i.p.), or when KET is directly applied to the intrathecal or intracranial ventricular space of rodents ^55–60^, KET-induced analgesia becomes OR dependent, and KET may also induce opioid-receptor dependent respiratory depression ^58^. This dose-dependent OR sensitivity is largely consistent with the low micromolar-range affinity of KET at the MOR ^7,46,61,62^, and suggests that at clinically relevant doses, KET’s opioid-receptor dependence is not mediated by direct agonism. Finally, we note that, in humans ^9^, NAL did not block the dissociative effects of KET, and dissociation may not be required for KET’s antidepressant effects ^63^. Similarly, when we tested dissociation using an assay of non-reflexive (“affective”) nociception (i.e. paw-licking during a hot plate test ^51^), KET’s effects were not MOR-dependent.

We found that NAL did not block KET’s effect in the FST and the TST, widely used to indicate antidepressant-like activity though criticized for their poor predictive value in identifying novel antidepressants ^36,38^. In addition to our present report, three independent laboratories working in mice and rats have found that NAL does not block KET’s ability to reverse immobility in the FST one day after treatment ^13,16,17^. Two groups have found that the *acute* KET effects in the FST are NAL-dependent ^16,17^, possibly reflecting the NAL-dependent locomotor stimulating effect which we report. These experiments, together with prior findings point to a reproducible model of MOR-dependent KET effects in rodents, characterized by a selective effect on locomotor activation and sensitization, but not analgesia, dissociation, or active coping assays.

### Brain-wide mapping highlights the CeA

Given the extensive literature on ketamine and other antidepressant effects on structural and functional plasticity in the hippocampus and prefrontal cortex ^21,64^, we were surprised that the CeA emerged from an unbiased, brain-wide mapping study of differential cFos expression comparing SAL+KET to NAL+KET. Follow-up imaging studies and experimental disruption of MOR signaling either pharmacologically or genetically confirm the role of MOR and PKCδ-expressing neurons of the CeA in locomotor activation and sensitization evoked by KET. Noting the well-described role of this neural population in acquired defensive behavior and salience detection ^31,65^, these behaviors may form the basis for future efforts at translational modeling.

CeA PKCδ neurons constitute a major population in the CeA that play critical roles in fear learning and extinction ^28,32^, states of anxiety, and hypophagia ^66,67^. Our locomotor data are consistent with the suggested role of these neurons in states of fear, anxiety, and behavioral inhibition. We identified *Oprm1* expression specifically in PKCδ neurons, further suggesting that NAL stimulates these neurons to disrupt KET’s effects on locomotion. These cells are also enriched with enkephalin, one of the primary endogenous agonists at MOR. Considering that PKCδ neurons project to limbic structures involved in the processing of affect and pain states, such as the bed nucleus of the stria terminalis, periaqueductal gray, and parabrachial nucleus, it will be important for future work to determine the precise interplay between ketamine and endogenous opioid signaling in the CeA.

### Limitations

We note two important limitations of this work: 1) CeA is a critical OR-sensitive node in a network that controls KET’s behavioral effects, but we have not identified how KET directly enhances locomotor behavior; and 2) we cannot distinguish between direct MOR-agonist effects of KET and indirect effects on endogenous opioids. Regarding the locomotor stimulating mechanism of KET, we draw on several strong prior studies showing KET enhances glutamatergic and dopaminergic transmission in the dorsal striatum, nucleus accumbens and PFC ^23,68,69^. These broad mechanisms have been noted for a range of drugs which, interestingly, all have abuse potential in humans and produce this simple behavior in mice ^42^. As we performed our brain mapping study with a threshold dose of KET (10 mg/kg i.p.), rather than a higher dose (20 mg/kg, Fig. 3h-j) which more reliably elicits locomotor sensitivity, we likely lacked the sensitivity to detect cFos changes in these other brain areas.

OR agonism and endogenous opioid release are among KET’s many pharmacological effects ^7,55,62^. However, our data strongly suggest that neither mechanism is relevant in the CeA. We replicated the effect of systemic NAL on KET-evoked hyperlocomotion by selectively deleting MORs from CeA neurons, and these neurons responded most robustly to NAL, rather than KET. As ORs are Gi/o-coupled, a direct effect at ORs in the CeA would be expected to produce a suppression of CeA activity, opposite to our results. This pattern of results is, again, most consistent with a permissive requirement for OR activation in KET’s behavioral effects. We cannot exclude the possibility that direct interactions with MORs or endogenous opioid release are also required elsewhere in the brain to produce KET’s locomotor stimulating effect.

### Future Directions

Developing novel antidepressant medications based on mechanistic insights has been exceptionally challenging given the very limited set of mechanism-informing experiments conducted in clinical populations. Conversely, classical behavioral pharmacology experiments like blocking or mimicking drug effects are relatively straightforward in animal models, but have rarely held up in human experiments. The novel in-human mechanistic finding that NAL blocks the antidepressant effect of KET offers the opportunity to develop new hypotheses about KET’s antidepressant mechanism and sets a new standard for evaluating translational models.

## ACKNOWLEDGEMENTS

We thank the Wu Tsai Neurosciences Institute Gene Vector and Virus Core for providing viruses. We thank the Wu Tsai Neuroscience Center, Neuroscience Microscopy Service for training and use of their LaVision BioTec UltraMicroscope II (S10OD025091-01). We thank John Sencaj for behavioral testing that was not included in the final version of the manuscript, Sofia R Schlozman and Dr. Tuuli M. Hietamies for helping with training Ilastik, and Dr. Vivianne Tawfik and her lab for assisting with behavioral analysis of pain assays. We also thank the entire Heifets and Malenka Labs for helpful discussions. This work was supported by NIH grants K99 DA056573 (M.B.P.), K08 MH110610, R01 MH130591 (B.D.H.), and P50 DA042012 (B.D.H. and R.C.M.), and grants from the Stanford University Wu Tsai Neurosciences Institute (R.C.M.).

## AUTHOR CONTRIBUTIONS

M.B.P., P.L., S.V., R.C.M., and B.D.H. conceived experiments and interpreted results. M.B.P. and B.D.H. wrote the manuscript, which was edited by all authors. S.V., and P.L. performed all behavioral assays and analyses. D.R.R., A.B.C., and J.S.S. performed iDISCO and UNRAVEL analysis and interpreted brain-wide mapping results. N.G. performed gait analysis. B.D.H. imaged the cleared brains. M.B.P. performed FISH. P.L. and S.V. performed immunohistochemistry.

## METHODS

### Subjects

Female and male C57BL/6J (Jackson Laboratory; stock #00664), heterozygous Fos-TRAP2 mice (Jackson Laboratory; stock #030323), heterozygous Gt(ROSA)26Sor^tm14(CAG-tdTomato)Hze^/J (Ai14, Jackson Laboratory; stock #007908), and homozygous *Oprm1*^tm1.1Cgrf^/KffJ (*Oprm1*^fl/fl^, Jackson Laboratory; stock #030074) mice were used. All mice (8-16 weeks old) were kept on a C57BL/6J background and group housed on a 12-hr light/dark cycle with food and water *ad libitum*. All procedures compiled with animal care standards set forth by the National Institute of Health and were approved by the Stanford University’s Administrative Panel on Laboratory Animal Care and Administrative Panel of Biosafety.

### Drug administration

Ketamine (10–30 mg/kg, MWI Veterinary Supply Co.), Naltrexone (5 mg/kg, Tocris, 0677), beta-funaltrexamine (β-FNA, 15 mg/kg, Sigma Aldrich, O003), norbinaltorphimine (norBNI, 10 mg/kg, Tocris, 0347), and cocaine (10 mg/kg, Sigma Aldrich, C5776) were all dissolved in 0.9% normal saline. 4-hydroxytamoxifen (4OHT, 50 mg/kg, Sigma Aldrich, T176) was dissolved in warm ethanol (50 µl / mg 4OHT) and mixed into sunflower oil (80 µl / mg 4OHT) and castor oil (20 µl / mg 4OHT), followed by vacuum centrifugation to evaporate out the ethanol. Drugs were administered intraperitoneally (i.p.) at a volume of 10 mL/kg. For all experiments involving drug injection, matched control groups always received the same total number of drug or SAL injections.

For local inhibition of MORs in the CeA, mice were bilaterally microinjected with CTAP (300 ng in 500 nL in saline, Sigma Aldrich, C6352) 5 min before behavioral testing. CTAP was infused through an injector cannula coupled to a 5 µL Hamilton syringe using a microinfusion pump (Harvard Apparatus) at a continuous rate of 150 nL min^-1^ to a total volume of 0.5 µL per hemisphere. Injector cannulas were removed 1 min after infusions were complete.

### Viral vectors and stereotaxic surgery

AAVs purchased from Stanford Neuroscience Gene Vector and Virus Core: AAVdj-hSyn-Cre-eGFP, AAVdj-hSyn-eGFP. Viruses were injected with 4-6 x 10^12^ infectious units per mL.

Mice of at least 7 weeks of age were anesthetized with isoflurane (1-2% v/v) and secured in a stereotaxic frame (David Kopf Instruments, Tujunga, CA). A syringe (Hamilton, Reno, Model 85 RN SYR) with a 33-gauge needle (VWR, no.7762) was used to infuse virus into the CeA (AP −1.20; ML ±2.95; DV −4.45) at a rate of 333 nL min^-1^ over 90 sec (500 nL total volume). Injector pipettes were slowly retracted after a 5 min diffusion period. The incision was then sealed with Vetbond (3M). For drug microinfusions, a 26-gauge bilateral cannula (P1 Technologies), 5.9 mm pitch, cut 3.7 mm in length from the cannula base, was implanted over the CeA (AP −1.34; ML ±2.95; DV −3.0). Cannulas were secured to the skull with stainless steel screws (thread size 00-90 x 1/16, Antrin Miniature Specialties), C&B Metabond, and light-cured dental adhesive cement (Geristore A&B paste, DenMat). Mice were group housed to recover for 3 weeks before experiments began.

### Inflammatory Pain Model

Mice received an intraplantar injection of Complete Freund’s adjuvant (CFA, 100 μl) in their right hind paw. Normal saline (0.9%) was used as a control. Testing was conducted 24 hrs after injection.

### Behavior

#### Open field test

Test mice were placed into the corner of an open field arena (40 x 40 cm) and allowed to move freely for a 30-min session. Time spent in the center (20 x 20 cm) was assayed as a measure of anxiety-like behavior. Total distance traveled and center time were measured automatically using BIOBSERVE. Mice were tested immediately after i.p. injections.

#### Forced Swim Test

The forced swim test (FST) was conducted in 5000 mL beakers filled with water kept at a temperature of 23C ± 0.5°C. No pre-test was conducted. Mice were administered drugs 24 hrs prior to starting the FST and tested for 6 min. Video analysis was completed blinded to treatments and manually scored. Only the last 4 min of the assay was scored.

#### Tail Suspension Test

The tail suspension test (TST) was conducted by hanging mice by their tails with a piece of tape and video recorded. Mice were administered drugs 24 hrs prior to starting the TST and tested for 6 min. Video analysis was completed blinded to treatment and manually scored. Only the last 4 min of the assay were scored.

#### von Frey

Analgesia was tested using the von Frey assay, with filaments ranging from 0.04 g – 6.0 g. Testing always began with a filament of 1.0 g using the up-down method for scoring. Animals were acclimated to individual chambers on a wire mesh for 30 min over 3 days before testing began. The filaments were applied to the left hind paw over 3 sessions, baseline, post-CFA injection and post-drug injection. The filaments were applied perpendicular to the hind paw and positive responses were recorded when the paw was lifted.

#### Hot Plate

The hot plate assay was used to assess dissociation through decreases in affective behaviors. Mice were habituated to the room for an hour before the start of testing. Mice were administered drugs or vehicle and placed on the hot plate (52.5° C) for 45 sec and video recorded. Analysis was conducted blinded to treatment.

#### Catwalk gait analysis

Mice were given drugs as above and then recorded freely ambulating on the Catwalk XT gait analysis system and were scored across 185 parameters (Noldus, Leesburg, VA, USA). A subset of critical parameters in each experiment was identified using random forest analysis, defined as a mean decrease in impurity > 0.02. This process identified 15 and 13 key parameters for the COC (Fig. S1e; Table S1) and KET+NAL (Fig. 1d,e; Table S2) experiments, respectively. These parameters were subsequently used in a two-component linear discriminant analysis (LDA). To assess the statistical significance of group differences based on the LDA projection, a multivariate analysis of variance (MANOVA) was employed. An ANOVA on the Euclidean distances between group centroids served as a post-hoc analysis to delineate which groups demonstrated statistically significant variances.

### Chronic Corticosterone

To study the effects of stress in mice, CORT was diluted in the drinking water for 21 days at a dose of 0.1 mg/ml. Mice were housed in groups of 4 and shared access to water. The solution was replaced with freshly diluted corticosterone every 3 days. Testing began after the 21 days of chronic corticosterone.

### Immunohistochemistry

For quantification of CeA cFos induction, mice were transcardially perfused with 4% PFA and postfixed overnight in the same solution. 24 hrs later, brains were transferred to phosphate buffered saline (PBS) and brains were cut coronally with a vibratome (50 μm), collecting alternating slices spanning the CeA. The free-floating sections were permeabilized with PBS– 0.3% Triton X-100 for 30 min, incubated for 1 hr at RT in a blocking solution with 5% goat serum in PBS–1% Triton X-100, and then incubated overnight at 4°C with a monoclonal cFos rabbit antibody (1:1000; 226 008; Synaptic Systems). Brain slices were rinsed in PBS three times for 10 min at RT, incubated for 1.5 hrs (RT) with donkey anti-rabbit Alexa 647 antibody (1:500; A32795; ThermoFisher Scientific) diluted in blocking solution, rinsed three times for 5 min, and mounted on slides with Fluoromount-G with DAPI (SouthernBiotech). Images were acquired on a Keyence BZ-X800 epifluorescence microscope using a 10x objective, 100% excitation, and no binning. Exposure times were 1/150 sec and 1/4 sec for DAPI and Cy5 filter cubes, respectively. Ilastik ^70^ was trained to segment cFos^+^ cells using all features and 12 training slices (3 slices / condition, with slices spanning the AP axis of the CeA). Offline analysis using ImageJ was performed for each brain slice in a uniform manner that was blind to mouse genotype. A region of interest (ROI) defining the left and right CeA for each slice was drawn to confine cFos+ cell counts and measure the area of the ROIs.

### Whole brain immunofluorescence (IF) staining and clearing

Mice were perfused with 1xPBS, followed by 4% PFA. Brains were dissected, post-fixed overnight, and washed thoroughly with 1xPBS. After adding 0.02% sodium azide, samples were stored at 4 °C until proceeding with IF labeling/iDISCO+, which followed published protocols ^22^.

### Whole brain imaging

After waiting more than 24 hrs for brains to clear in dibenzyl ether, they were imaged using a LaVision BioTec UltraMicroscope II with a 2x detection objective and 0.8x zoom, yielding x and y voxel dimensions of 3.78 µm. At this magnification, the entire sample was contained in the field of view. Cleared hemispheres were mounted using a c-clamp, oriented with the medial side down for sagittal images. Dibenzyl ether was used as the imaging solution (refractive index: 1.561). Images were acquired using ImspectorPro software. The instrument mode was set to CDC 2x. The light sheet width was set to 100% and thickness was set to 5 µm. The sheet NA was 0.156. Laser power was 20-24% for 488 nm and 100% for 640 nm. The exposure time was 25 ms for 488 nm and 100 ms for 640 nm. Left and right light sheets were aligned. The horizontal focus was set at two locations in the light sheet plane. Left and right light sheets were merged using a blending algorithm. A series of images were acquired spanning each hemisphere with a z-step of 2.5 µm and saved as 16-bit .ome.tifs. X and y images dimensions were 2160 and 2560 pixels, respectively.

### Image pre-processing

Image pre-processing, voxel-wise statistics, and cluster validation were automated with UNRAVEL (Rijsketic, Casey et al., 2023). In brief, autofluorescence image volumes were thresholded by adjusting the display range to zero-out external voxels, 2x downsampled, reoriented, and converted to .nii.gz images. In cases where the hemisection was not perfectly down the midline, autofluorescence images were fixed for registration in 3D Slicer by digitally trimming excess tissue or adding missing tissue from the contralateral hemisphere. Additionally, fake tissue was added to account for occlusion from the c-clamp and/or tissue damage. These modifications improve intensity matching between the autofluorescence images and the average template brain during registration. Likewise, we used an iDISCO/LSFM-specific version ^24^ of the Allen Mouse Brain Common Coordinate Framework (CCFv3; REF) for registration. Prior to analysis, data was omitted from three hemispheres immunostained for cFos due to missing raw data, a large bubble casting a shadow through the tissue, or tissue damage combined with poor registration accuracy. cFos immunofluorescence (cFos-IF) was rolling ball background subtracted using a pixel radius of 4. Transformation matrices from registration were used to warp background subtracted cFos-IF images to atlas space. Intensities were normalized by z-scoring each hemisphere, using a brain mask lacking the dissected olfactory bulbs.

### Voxel-wise statistics

Similar to our prior report ^22^ voxel-wise statistics were performed using FSL (FMRIB Software Library 6.0.2:a4f562d9; www.fmrib.ox.ac.uk/fsl) to identify clusters of differential cFos-IF. Briefly, background subtracted, z-scored, cFos-IF image volumes in 25 μm LFSM atlas space were merged into a 4D image file using fslmerge and smoothed with a 50 μm full width at half maximum Gaussian filter. The randomise_parallel tool in FSL was used to carry out a two-sample unpaired t-test comparing group differences to a null distribution based on nonparametric permutation testing (90,000 permutations) with the general linear model. Multiple comparisons correction of voxel-wise statistical contrasts was performed using an FDR threshold of q < 0.05 (uncorrected p ≤ 0.0000075) and a cluster extent threshold of 100 voxels (≥1.156 x 10^-3^ mm^3^). A hemispheric mask containing only voxels corresponding to grey matter present in all samples defined voxels to be analyzed (n = 6,252,542 voxels across 493 brain regions). Tissue masks were generated by thresholding each cFos-IF image in atlas space to zero-out voxels lacking apparent tissue, followed by binarization, summation of all tissue masks (n=15), thresholding with an intensity of 15, and binarization. Voxels corresponding to fiber tracts, ventricles, or excised olfactory bulbs were excluded by making a mask of the atlas that omitted these regions. This mask was applied to the group-level tissue mask to create the mask used for FDR correction.

### Cluster validation

Clusters surviving FDR correction and extent thresholding were warped to tissue space for each sample. Using numpy, each cluster was cropped, measured volumetrically, and converted to a binary mask. Cluster-specific masks in tissue space were then applied to consensus cFos or tdTomato segmentation images generated using the Ilastik pixel classification workflow, as previously described ^22^. Custom Fiji macros performed object counting, which, in combination with cluster volumes, were used to calculate the cell density within each cluster for all samples. *Post hoc* t-tests were performed in GraphPad Prism 10.

### Fluorescence in situ hybridization

For examination of gene expression in the CeA, wild-type brains were flash frozen in isopentane on dry ice, sectioned on a cryostat at 16 µm, and processed for fluorescence *in situ* hybridization (FISH) with the RNAscope Multiplex Fluorescent v2 assay according to the manufacturer’s guidelines (Advanced Cell Diagnostics). Transcripts examined were *Fos* (ACDBio cat# 316921-C2), *Oprm1* (ACDBio cat# 315841-C3), *Pkcd* (ACDBio cat# 441791), and *Sst* (ACDBio cat# 404631). Slides were coverslipped with Fluoromount-G with DAPI (Southern Biotech, 0100-20) and stored at 4°C in the dark before imaging. Application of protease III for 5 min was used as opposed to protease IV for 30 min. Fluorescent images were collected on a Nikon A1 confocal microscope. Each image was acquired using identical pinhole, gain, offset, and laser settings (1024 × 1024 pixels). Cells were counted and co-localization was measured using Fiji (Schindelin et al., 2012). The number of *Fos* cells expressing *Oprm1*, *Pckd*, and *Sst* was calculated within each CeA section and averaged for each subject.

## QUANTIFICATION AND STATISTICAL ANALYSIS

All behavioral data were analyzed and graphed with GraphPad Prism (9 or 10). Data distribution and variance were tested using Shapiro-Wilk normality tests. Normally distributed data were analyzed by unpaired, two-tailed t-tests, or one- or two-factor repeated measures ANOVA with *post-hoc* Sidak or Tukey correction. When normal distributions were not assumed, the Wilcoxon signed rank test was performed for within group comparisons of two treatments, Mann-Whitney for between group comparisons, and Kruskal-Wallis with post-hoc Dunn’s test for multiple comparisons. No statistical methods were used to predetermine sample size, which was based on extensive prior experience with the assays used. All experiments were conducted in a blinded manner such that assays were conducted and analyzed without knowledge of the specific manipulation being performed and with animals being randomized by cage before behavioral experiments. Differences were considered significant when p < 0.05. All pooled data are expressed as mean ± SEM.

## SUPPLEMENTAL FIGURE LEGENDS

**Figure S1.**
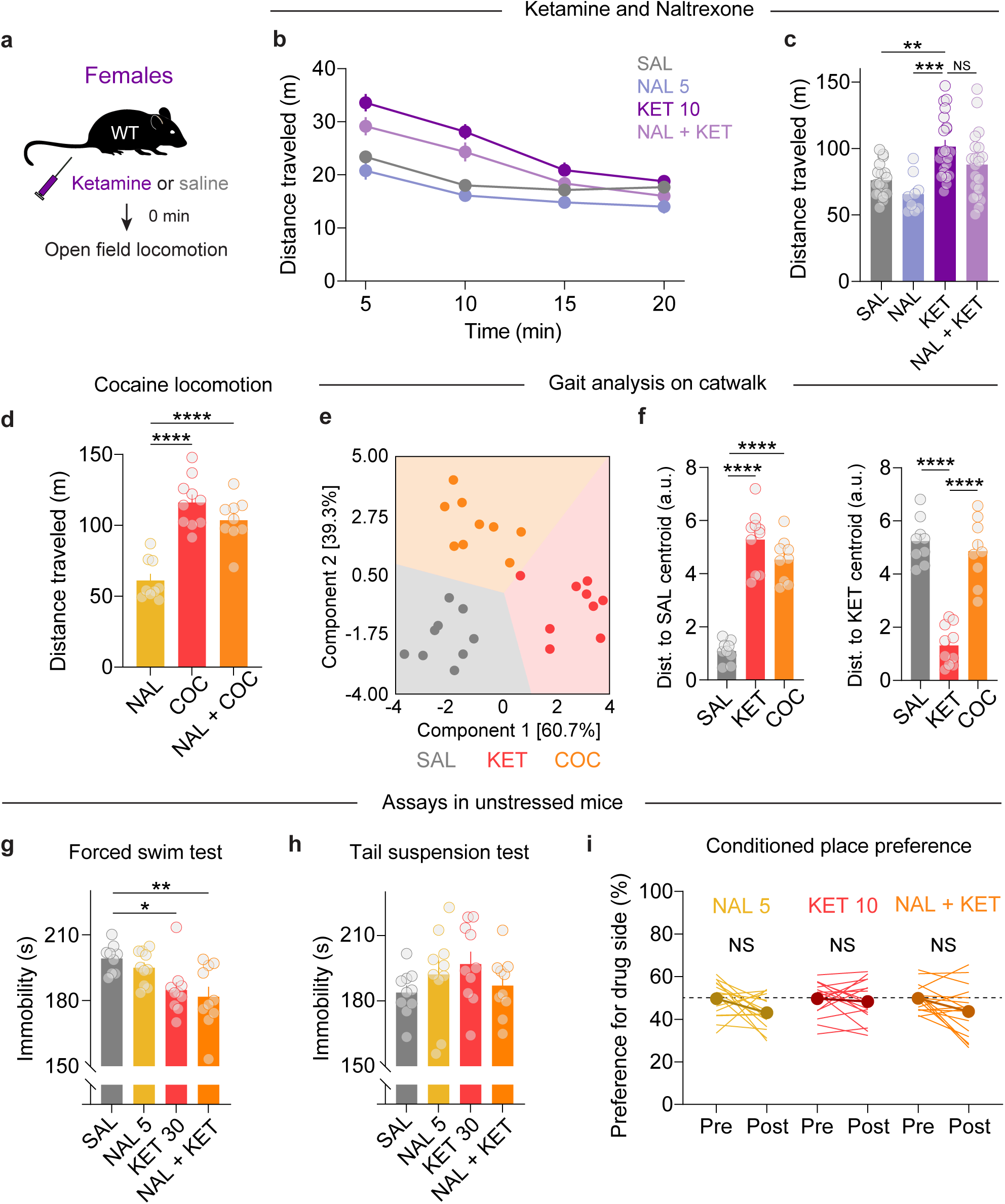
Ketamine locomotor effects in females and additional behavioral metrics. a) Schematic of experimental procedure and time points in female mice for panels b-c. b) Time course of locomotor activity after SAL (n=17), NAL (5 mg/kg; n=9), KET (10 mg/kg; n=21), or NAL+KET (n=21). c) Summary of total distance traveled in the open field test. KET increased total distance traveled, an effect largely unaltered by naltrexone. Ordinary one-way ANOVA, F_3,64_=8.17, ***p=0.0001. Tukey’s multiple comparisons test, KET vs SAL **p = 0.0021, KET vs NAL ***p = 0.0003, KET vs NAL + KET p = 0.15. d) In male mice, COC (15 mg/kg) locomotion was not blocked by NAL (5 mg/kg). NAL, n=9; COC, n=10; NAL+COC, n=9. Ordinary one-way ANOVA, F_2,25_ = 29.37, ****p < 0.0001. Tukey’s multiple comparisons test, KET vs COC ****p < 0.0001, KET vs NAL + COC ****p < 0.0001. e) Two-component linear discriminant analysis (LDA) of mouse gait after SAL (n=9), KET (10 mg/kg; n=10) or COC (15 mg/kg; n=9). See Table S2 for component 1 and 2 feature weights. Colored fields represent LDA decision boundaries. All subjects were accurately classified. A significant drug effect was observed. MANOVA, Pillai’s trace = 0.78, F_2,25_ = 43.33, p <0.001. f) Distance (a.u.) of all groups to either SAL (*left*) or KET (*right*) centroid. KET subjects are a significant distance from both SAL and COC centroids. ANOVA, SAL centroid F_2,25_ = 64.38, p < 0.0001. Tukey’s multiple comparisons test, KET vs SAL **** p < 0.0001, COC vs SAL **** p < 0.0001, KET vs COC p = 0.15. ANOVA, KET Centroid F_2,25_ = 50.30, p < 0.0001. Tukey’s multiple comparison test, SAL vs KET **** p < 0.0001, COC vs KET **** p < 0.0001, SAL vs COC p = 0.68). g) KET (30 mg/kg) decreased immobility in the FST in unstressed mice. NAL did not reverse this effect. SAL, n=9; NAL, n=10; KET, n=10; NAL+KET, n=10. One way ANOVA, F_3,35_ = 5.82, **p = 0.0025. Tukey’s multiple comparisons test, SAL vs KET 30 *p = 0.026, SAL vs NAL + KET **p = 0.0053. h) KET did not alter immobility in the TST in unstressed mice. SAL, n=10; NAL, n=10; KET, n=10; NAL+KET, n=10. One way ANOVA, F_3,36_ = 1.26, p = 0.30. i) Unstressed mice were tested for conditioned place preference to NAL (n=16), KET (20 mg/kg; n=16), or NAL+KET (n=16). 2-way RM ANOVA, interaction of drug and time, F_2,45_ = 1.04, p = 0.36.

**Figure S2.**
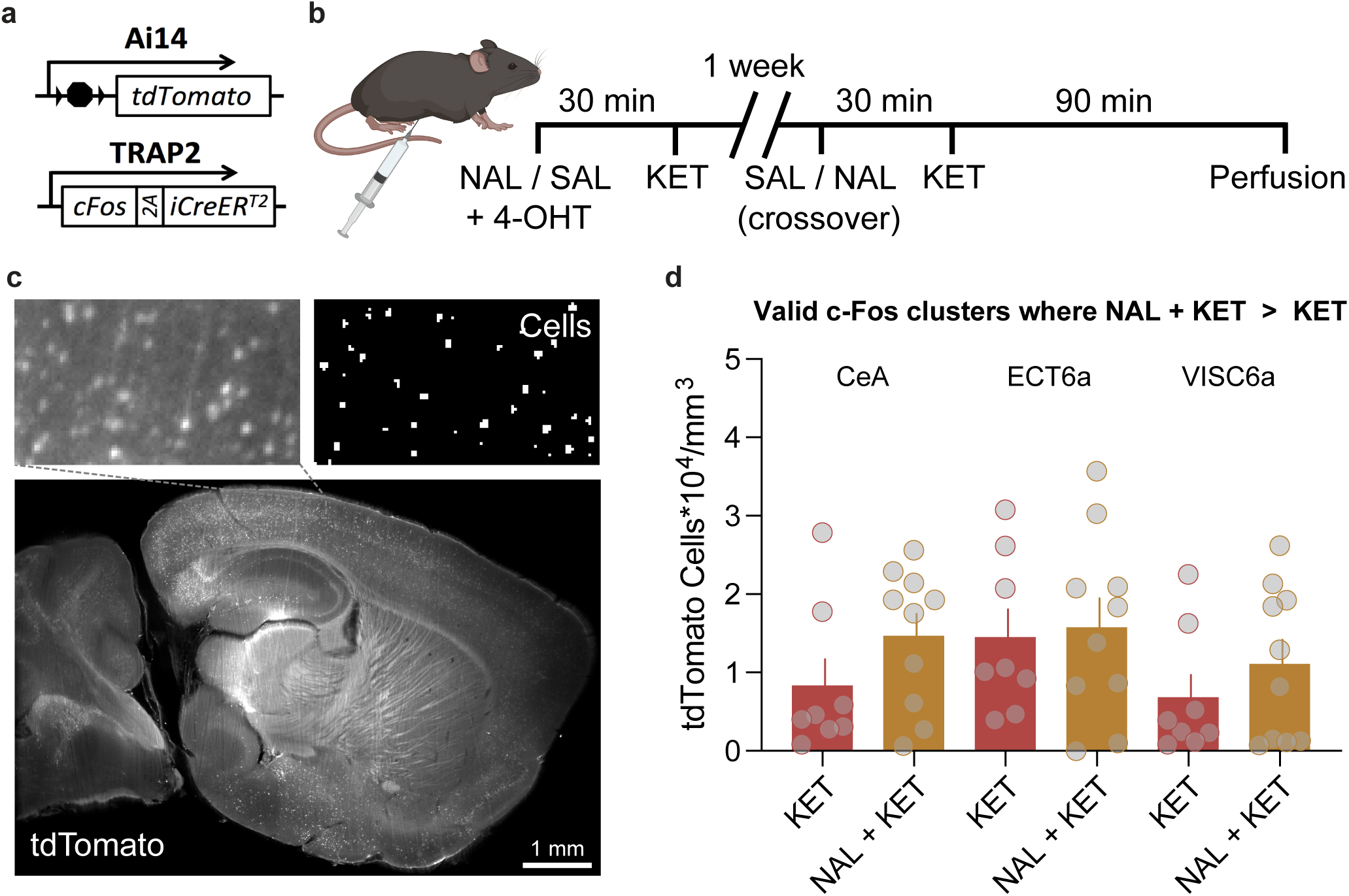
Whole brain mapping in TRAP2;Ai14 mice. a) Schematic illustrating the TRAP2;Ai14 expression system. b) Timeline of experimental procedure. c) Example image of tdTomato expression in an iDISCO-cleared brain. d) NAL+KET (n=10) increased tdTomato^+^ cell densities relative to KET (n=8) in the same regions cFos was detected as increased, albeit less robustly. Unpaired 2-tailed t-tests were performed for each cluster: CeA, p = 0.16; ECT6a, p = 0.81; VISC6a, p = 0.34.

**Figure S3.**
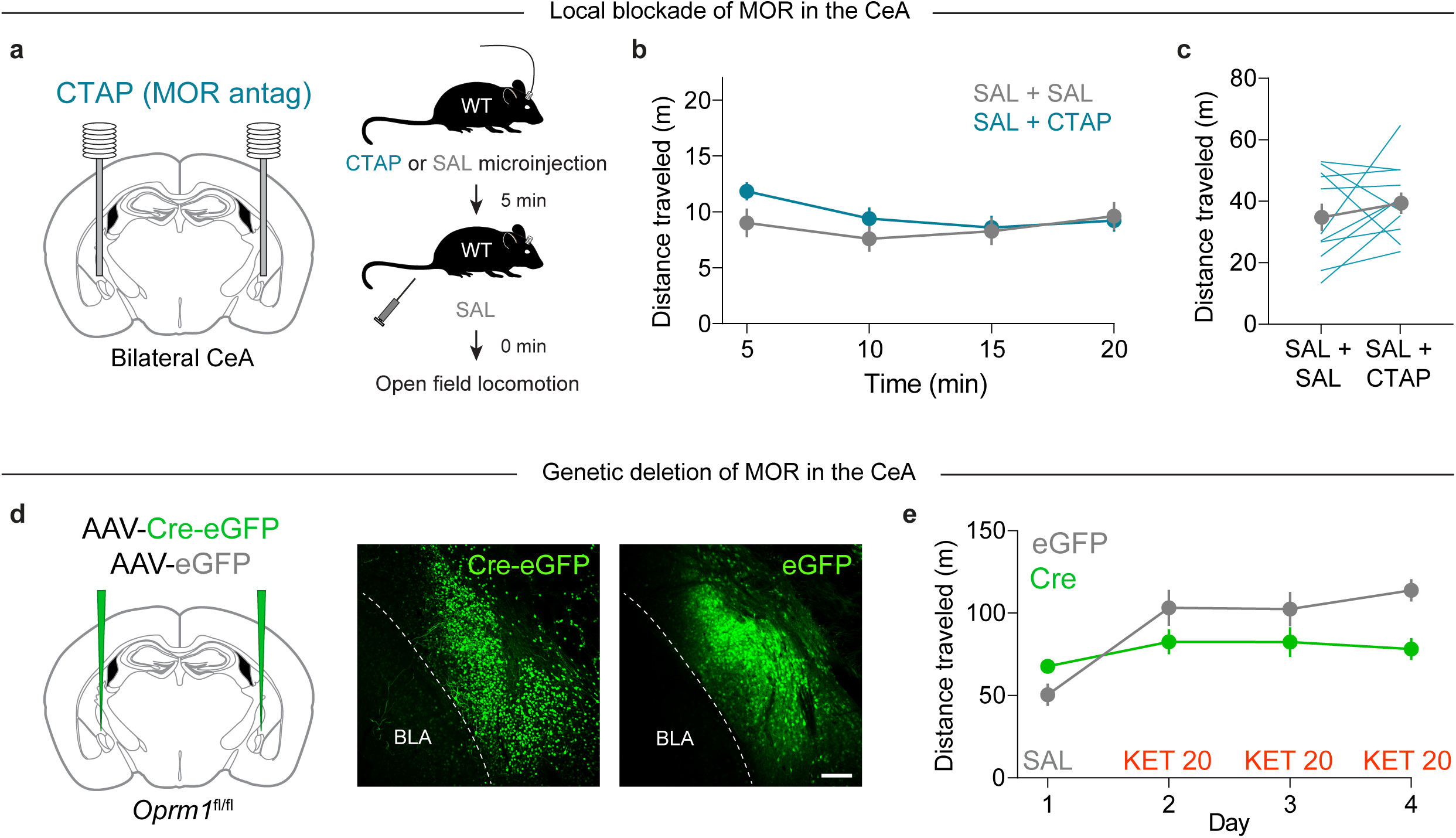
MORs in CeA are critical to ketamine’s locomotor effects. a) Left, setup of MOR antagonist microinjection into CeA before a control saline injection. Right, experimental procedure. b) Time course of locomotor activity after administration of saline and CTAP microinjection (300 ng in 500 nL / hemisphere). SAL+SAL, n=11, SAL+CTAP, n=11; crossover design. c) CTAP had no effect on baseline locomotor behavior in the open field. Paired, two-tailed t-test, t_10_ = 1.13, p = 0.29. d) Left, injection of AAV-Cre-eGFP into the CeA of *Oprm1*^fl/fl^ mice for genetic deletion of MOR. Right, representative images of Cre-eGFP and eGFP expression in the CeA. Scale = 100 µm. e) Time course of locomotor sensitization to KET (20 mg/kg) in mice with genetic deletion of MOR in the CeA (Cre, n=14) or a control group (eGFP, n=5). This is the same data presented in Fig. 3e, now without normalization to Day 1 (SAL). 2-way RM ANOVA, interaction of *Oprm1* cKO and drug condition, F_3,51_ = 9.73, ****p < 0.0001. Sidak’s multiple comparisons test, eGFP vs Cre: Day 1 **p = 0.0011, Day 2 p=0.67, Day 3 p = 0.35, Day 4 **p < 0.0018.

**Table S1.** Summary of feature importance, weights and component dimension scaling factors for COC, KET, SAL catwalk analysis.

| Feature | Mean Decrease<br>in Impurity | Feature Weights |  |  | Scaling for Fig. S1e |  |
| --- | --- | --- | --- | --- | --- | --- |
|  |  | Saline | Ketamine | Cocaine | Component 1 | Component 2 |
| Right Front Paw Step Cycle Mean Duration (seconds) | 0.0482 | -12.52 | 9.50 | 1.96 | 3.76 | 2.39 |
| Coupling of Right Front Paw to Right Hind Paw Cstat Mean | 0.0362 | -27.30 | 31.81 | -8.04 | 11.44 | 1.43 |
| Bodyspeed During Right Hindlimb Contact | 0.0291 | -12.11 | 17.73 | -7.59 | 6.13 | -0.60 |
| Time to Cross Catwalk | 0.0291 | 12.87 | 17.50 | -32.32 | 4.10 | -11.75 |
| Phase Dispersion Right Front Paw to Right Hind Paw | 0.0282 | -29.71 | 13.79 | 14.38 | 6.37 | 8.65 |
| Right Hindlimb Swing Speed | 0.0267 | 2.31 | 2.43 | -5.00 | 0.53 | -1.86 |
| Step Cadence | 0.0257 | -10.06 | 5.75 | 3.67 | 2.47 | 2.56 |
| Average Speed | 0.0255 | 0.55 | -3.63 | 3.48 | -1.10 | 0.99 |
| Right Front Limb Stride Length Mean | 0.0252 | 5.51 | 0.00 | -5.52 | -0.43 | -2.48 |
| Phase Dispersions Right Front to Right Hind Cstat Mean | 0.0237 | 66.11 | -53.60 | -6.55 | -20.86 | -11.44 |
| Left Front Step Cycle Mean Duration (seconds) | 0.0235 | -0.36 | -21.70 | 24.47 | -6.31 | 7.55 |
| Print Positions of Right Paws | 0.0224 | 0.28 | 0.12 | -0.41 | 0.01 | -0.16 |
| Left Front Limb Stride Length | 0.0208 | -7.46 | 8.01 | -1.44 | 2.93 | 0.62 |
| Bodyspeed During Left Front Limb Contact | 0.0207 | 25.50 | -28.19 | 5.82 | -10.24 | -1.85 |
| Coupling of Right Front Paw to Right Hind Paw | 0.0201 | -16.38 | 11.03 | 4.12 | 4.51 | 3.60 |

**Table S2.** Summary of feature importance, weights and component dimension scaling factors for SAL/SAL, SAL/KET, NAL/KET catwalk analysis.

| Feature | Mean Decrease in Impurity | Feature Weights |  |  | Scaling for Fig. S1e |  |
| --- | --- | --- | --- | --- | --- | --- |
|  |  | SAL/SAL | SAL/KET | NAL/KET | Component 1 | Component 2 |
| Average Run Speed | 0.0460 | -2.21 | 5.96 | -3.75 | 2.56 | -2.93 |
| Duration of Catwalk Crossing | 0.0353 | 0.10 | 0.47 | -0.58 | 0.17 | -0.61 |
| Left Front Paw Print Area | 0.0353 | 1.67 | 1.56 | -3.23 | 0.36 | -3.86 |
| Left Hind Paw Ground Contact Maximum Intensity | 0.0345 | -1.06 | 1.90 | -0.84 | 0.87 | -0.44 |
| Coupling Between Left Front and Right Hind Limbs | 0.0327 | 1.40 | -1.25 | -0.15 | -0.66 | -0.67 |
| Left Front Paw Print Width | 0.0322 | 1.81 | -4.11 | 2.30 | -1.80 | 1.63 |
| Right Hindlimb Average Contact Intensity | 0.0318 | 1.59 | -1.51 | -0.09 | -0.79 | -0.68 |
| Body Speed During Right Front Paw Contact | 0.0311 | 2.55 | 1.34 | -3.88 | 0.16 | -4.84 |
| Average Catwalk Crossing Speed | 0.0275 | 4.31 | -17.59 | 13.28 | -7.25 | 11.69 |
| Left Front Paw Maximum Contact Area | 0.0234 | -2.37 | 0.37 | 1.99 | 0.46 | 2.88 |
| Left Front Limb Swing Speed | 0.0225 | -0.63 | 1.49 | -0.86 | 0.65 | -0.63 |
| Timing of Left Hind Limb Max Contact Area | 0.0206 | 0.72 | -0.83 | 0.11 | -0.41 | -0.16 |
| Body Speed During Left Hind Paw Contact | 0.0204 | -4.38 | 10.71 | -6.33 | 4.66 | -4.70 |

**Table S3.** Volume, location and composition of validated clusters of cFos immunofluoresence, KET vs NAL+KET.

| Table S3. Volume, location and composition of validated clusters of cFos immunofluorescence, KET vs NAL+KET |  |  |  |  |  |  |
| --- | --- | --- | --- | --- | --- | --- |
| Cluster | p value | Volume | CoG | Region name | Abbreviation | % of cluster volume |
| 1 | <0.00001 | 0.02177 | 283,265,71.8 | Central amygdalar nucleus, capsular part | CEAc | 77.67 |
|  |  |  |  | Basolateral amygdalar nucleus, anterior part | BLAa | 8.04 |
|  |  |  |  | Central amygdalar nucleus, lateral part | CEAl | 7.18 |
|  |  |  |  | Striatum | STR | 3.02 |
|  |  |  |  | Central amygdalar nucleus, medial part | CEAm | 2.94 |
|  |  |  |  | Intercalated amygdalar nucleus | IA | 1.01 |
| 4 | 0.00034 | 0.00328 | 329,306,115 | Ectorhinal area/Layer 6a | ECT6a | 100.00 |
| 5 | 0.49978 | 0.00281 | 323,256,115 | Visceral area, layer 6a | VISC6a | 93.33 |
|  |  |  |  | Visceral area, layer 5 | VISC5 | 4.44 |
|  |  |  |  | Supplemental somatosensory area, layer 6a | SSs6a | 2.22 |

